# Adolescent chronic sleep restriction promotes alcohol drinking in adulthood: evidence from epidemiological and preclinical data

**DOI:** 10.1101/2023.10.11.561858

**Authors:** Oluwatomisin O. Faniyan, Daniele Marcotulli, Reyila Simayi, Federico Del Gallo, Sara De Carlo, Eleonora Ficiarà, Doretta Caramaschi, Rebecca Richmond, Daniela Franchini, Michele Bellesi, Roberto Ciccocioppo, Luisa de Vivo

## Abstract

Epidemiological investigations have indicated that insufficient sleep is prevalent among adolescents, posing a globally underestimated health risk. Sleep fragmentation and sleep loss during adolescence have been linked to concurrent emotional dysregulation and an increase in impulsive, risk-taking behaviors, including a higher likelihood of substance abuse. Among the most widely used substances, alcohol stands as the primary risk factor for deaths and disability among individuals aged 15-49 worldwide. While the association between sleep loss and alcohol consumption during adolescence is well documented, the extent to which prior exposure to sleep loss in adolescence contributes to heightened alcohol use later in adulthood remains less clearly delineated.

Here, we analyzed longitudinal epidemiological data spanning 9 years, from adolescence to adulthood, including 5497 participants of the Avon Longitudinal Study of Parents And Children cohort. Sleep and alcohol measures collected from interviews and questionnaires at 15 and 24 years of age were analyzed with multivariable linear regression and a cross-lagged autoregressive path model. Additionally, we employed a controlled preclinical experimental setting to investigate the causal relationship underlying the associations found in the human study and to assess comorbid behavioral alterations. Preclinical data were collected by sleep restricting Marchigian Sardinian alcohol preferring rats (msP, n=40) during adolescence and measuring voluntary alcohol drinking concurrently and in adulthood. Polysomnography was used to validate the efficacy of the sleep restriction procedure. Behavioral tests were used to assess anxiety, risky behavior, and despair.

In humans, after adjusting for covariates, we found a cross-sectional association between all sleep parameters and alcohol consumption at 15 years of age but not at 24 years. Notably, alcohol consumption (Alcohol Use Disorder Identification Test for Consumption) at 24 years was predicted by insufficient sleep at 15 years whilst alcohol drinking at 15 years could not predict sleep problems at 24. In msP rats, adolescent chronic sleep restriction escalated alcohol consumption and led to increased propensity for risk-taking behavior in adolescence and adulthood.

Our findings demonstrate that adolescent insufficient sleep causally contributes to higher adult alcohol consumption, potentially by promoting risky behavior.

## Introduction

Epidemiological research has shown that adolescents worldwide are chronically sleep deprived due to a combination of physiological, social, and environmental factors [1] which reduce sleep duration and quality across adolescence [2–4]. Increased use of technology at night [5], consumption of caffeine, academic and social demands combine with ontogenetic preference towards later sleep and wake up times and lead to reduced sleep duration and quality across adolescence [2–4]. Sleep fragmentation and sleep loss have been associated with emotional dysregulation [6, 7], increased psychosis [8, 9], and higher impulsive and risk-taking behaviors [10, 11], including higher substance abuse [12, 13].

Recent reports show that alcohol remains the substance most widely used by today’s teenagers and represents the top risk factor for deaths and disability among the world’s population aged 15-49 years [14, 15]. Although genetic predisposition is estimated to contribute to approximately 50– 60% of the vulnerability to develop Alcohol Use Disorders (AUD) [16, 17], environmental factors such as sleep patterns, can also play an important role, especially during adolescence.

Cross-sectional studies showed a bidirectional association between sleep and circadian disturbances and alcohol problems [18–20]. Moreover, studies of communities and clinical samples found associations between different sleep difficulties (e.g. troubles falling or staying asleep, insomnia, restless sleep) in late childhood and adolescence and different aspects of alcohol use, including age of first alcohol use and AUD onset in adolescence and young adulthood [21–25]. In substance-naive 12 year old adolescents, shorter sleep and greater daytime sleepiness predicted the onset of alcohol use, heavy drinking, and alcohol-related consequences over a 4-year period [26]. These studies support the hypothesis that sleep impairment promotes alcohol use during adolescence, but the observation of this association is confined mostly to relatively short time intervals within adolescence. Whether sleep disruption during adolescence leads to higher alcohol consumption in adulthood and increases the risk of developing AUD and other psychiatric disorders later in life is unclear.

Here, drawing on available longitudinal data of sleep and alcohol drinking at two time points 9 years apart, in mid adolescence and young adulthood, we examined the cross-sectional and longitudinal associations between insufficient sleep and alcohol use in a large prospective cohort study of adolescents in the United Kingdom. We aimed at determining that self-reported insufficient sleep at 15 years of age was associated with higher alcohol use at the same age and that it was predictive of higher alcohol consumption at 24 years in this population.

Since epidemiological studies are correlative in nature, to test the direct causal link between exposure and outcome variables, tightly controlled experimental settings using animal models are crucial. Preclinical investigations can rigorously control for genetic, psychological, and environmental factors that could contribute to the development of AUD and neuropsychiatric comorbidities and potentially confound the associations found in human studies.

Preclinical experiments manipulating sleep during adolescence and looking at the subsequent effects on alcohol drinking in adulthood are lacking. Hence, we tested the longitudinal causal link between adolescent insufficient sleep and voluntary alcohol consumption using a preclinical model of chronic sleep restriction in a line of genetically selected alcohol preferring rats with innate predisposition to drink alcohol. Doing so, allowed us to track the evolution of voluntary alcohol consumption from early adolescence to adulthood. Additionally, this investigation used behavioral tests to measure the emergence of neuropsychiatric comorbidities in late adolescence and adulthood.

## Methods

### Epidemiological study

#### Cohort study numbers

The Avon Longitudinal Study of Parents And Children (ALSPAC) is a population-based prospective cohort situated in South West England. Detailed information about the methods and procedures of ALSPAC study is available elsewhere [27–30]. Briefly, pregnant women resident in Avon, UK, with expected dates of delivery between 1st April 1991 and 31st December 1992 were invited to take part in the study. 20,248 pregnancies have been identified as being eligible and the initial number of pregnancies enrolled was 14,541. Of the initial pregnancies, there was a total of 14,676 foetuses, resulting in 14,062 live births and 13,988 children who were alive at 1 year of age. When the oldest children were approximately 7 years of age, an attempt was made to bolster the initial sample with eligible cases who had failed to join the study originally. As a result, when considering variables collected from the age of seven onwards there are data available for more than the 14,541 pregnancies mentioned above: The number of new pregnancies not in the initial sample (Phase I enrolment) that are currently represented in the released data and reflecting enrolment status at the age of 24 is 906, resulting in an additional 913 children being enrolled (456, 262 and 195 recruited during Phases II, III and IV respectively). The total sample size for analyses using any data collected after the age of seven is therefore 15,447 pregnancies, resulting in 15,658 foetuses. Of these, 14,901 children were alive at 1 year of age. All participants have been followed up regularly with questionnaires on lifestyle behaviors and research clinics.

The study website contains details of all the data that is available through a fully searchable data dictionary and variable search tool (http://www.bristol.ac.uk/alspac/researchers/our-data/). Ethical approval for the study was obtained from the ALSPAC Ethics and Law Committee and the Local Research Ethics Committees.

#### Participants

The present study included only the participants who attended the teen focus clinic at 15 years (n=5497), between October 2006 and November 2008. At the latter time point of our analyses, participants were 24-year-old. Study data were collected and managed using REDCap electronic data capture tools hosted at the University of Bristol [31]. REDCap (Research Electronic Data Capture) is a secure, web-based software platform designed to support data capture for research studies. Informed consent for the use of data collected via questionnaires and clinics was obtained from participants following the recommendations of the ALSPAC Ethics and Law Committee at the time.

#### Exposure: Sleep

At 15 years, sleep was measured with the School Sleep Habits Survey [32] and sleep parameters were derived from the following items: “Assessment of how much sleep the participant gets” (3-point Likert type scale: 1=too much, 2=enough, 3=too little) for *sleep quantity*; “Length of time the participant sleeps on normal school night” for *hours of sleep per night*; “Problem participant has with sleepiness during daytime activities” for *daytime sleepiness* (5-point Likert type scale: 1=none, 2=a little, 3=more than a little, 4=big problem, 5=very big problem), and “Frequency participant feels that they get enough sleep” (5-point Likert type scale: 1=always, 2=usually, 3=sometimes, 4=rarely, 5=never) for *sleep satisfaction* [33]. We computed a *sleep insufficiency score* (min=0, max =3) at 15 years by summing up the minmax scaled scores of the questions for sleep quantity, daytime sleepiness, and sleep satisfaction to summarize both the quantity and quality domains of sleep.

At 24 years, participants completed the Health and Wellbeing CIS-R questionnaire [34]. Sleep measures were derived from the following questions: “In past month, participant had problems getting to sleep or back to sleep” (1=no; 2=yes); “In past 7 nights, number of nights participant had sleep problems” (1=no; 2= between one and three nights; 3=four nights or more); “In past week, length of time participant got to sleep on night with least sleep” (1=less than 15 min; 2=between 15 minutes and 1 hour; 3=between 1 and 3 hours; 4=three hours or more); “In past 7 days, number of nights participant spent >=3 hours getting to sleep”; “In past 7 days, participant woke >2 hours early & couldn’t get back to sleep” (1=none; 2=between one and three nights; 3=four nights or more). We then constructed a total sleep insufficiency score for this age by summing up the minmax scaled scores of the five questions.

#### Outcome: Alcohol consumption

Alcohol consumption at 15 years was calculated by multiplying the frequency at which the child reported to drink alcohol by the number of drinks consumed on a normal drinking day, as in [35]. The presence and severity of alcohol use disorder at 24 years was investigated using the Alcohol Use Disorder Identification Test for Consumption (AUDIT-C) [36, 37], obtained in a computer-assisted clinical interview.

#### Covariates

Based on previous literature of confounding factors associated with both exposure and outcome variables [38–41], biological sex at birth, mother’s AUDIT, mother’s occupational class, used as a proxy for maternal socioeconomic status, and presence of any psychiatric disorder at 15 years were included as covariates. Mother’s occupational class was derived at 32 weeks of gestation and used as a proxy for maternal socioeconomic status using the 1991 Office of Population, Censuses and Surveys classification in UK (I-professional occupations, II-managerial and technical occupations, III skilled non-manual occupations, III skilled manual occupation, IV-partly skilled occupation, V-unskilled occupation) [42]. Mother’s AUDIT-C was completed when the participant was 18 years old. The presence of psychiatric disorders was derived from the questionnaire Development and Well-Being Assessment (DAWBA) completed by the participant at 15 years of age, which assesses the likelihood of any psychiatric disorder (emotional, behavioural, and hyperactivity disorders) in one of six bands [43, 44].

#### Missing data

Table 1 describes the percentage of missing values for each variable. Attrition is a well-known problem in the ALSPAC dataset like other longitudinal studies and a complete-cases analysis could result in potentially biased findings [45]. Therefore, we imputed missing data. Univariate logistic regressions were used to evaluate the association between the independent variables and outcome data missingness. We observed that children from lower socioeconomic backgrounds, males, and individuals with mental health issues were more likely to skip follow-ups (Table 1), which indicates a relationship between the observed variables and data missingness. For this reason, exclusively including participants with complete data might potentially result in selection bias. To address this, we employed multivariable imputation by chained equations to impute missing data [46] using the ‘futuremice’ command in the ‘mice’ R package [47]. We imputed a total of 50 complete datasets (n = 5497 all participants at the 15 years focus clinic) through 5 imputation iterations. The imputation matrix included all the variables in the regression models (biological sex at birth, alcohol consumption at 15 years, AUDIT-C at 24 years, mother’s AUDIT, mother’s social class, sleep measures at 15 and 24 years), and an additional set of auxiliary variables related to sleep issues, alcohol consumption, socioeconomic status, adverse childhood experiences, and adolescent psychopathology at 15 years, 24 years, and at multiple time points before 24 years. Including this extra set of auxiliary variables increased the accuracy of the imputation by providing more information to estimate the missing data, and supported the assumption that data missingness was conditional on the factors we could observe and include in the imputation model (such as a participant’s socioeconomic status or sex) rather than on the ones we couldn’t include (Missing at Random assumption) [48]

**Table 1.**
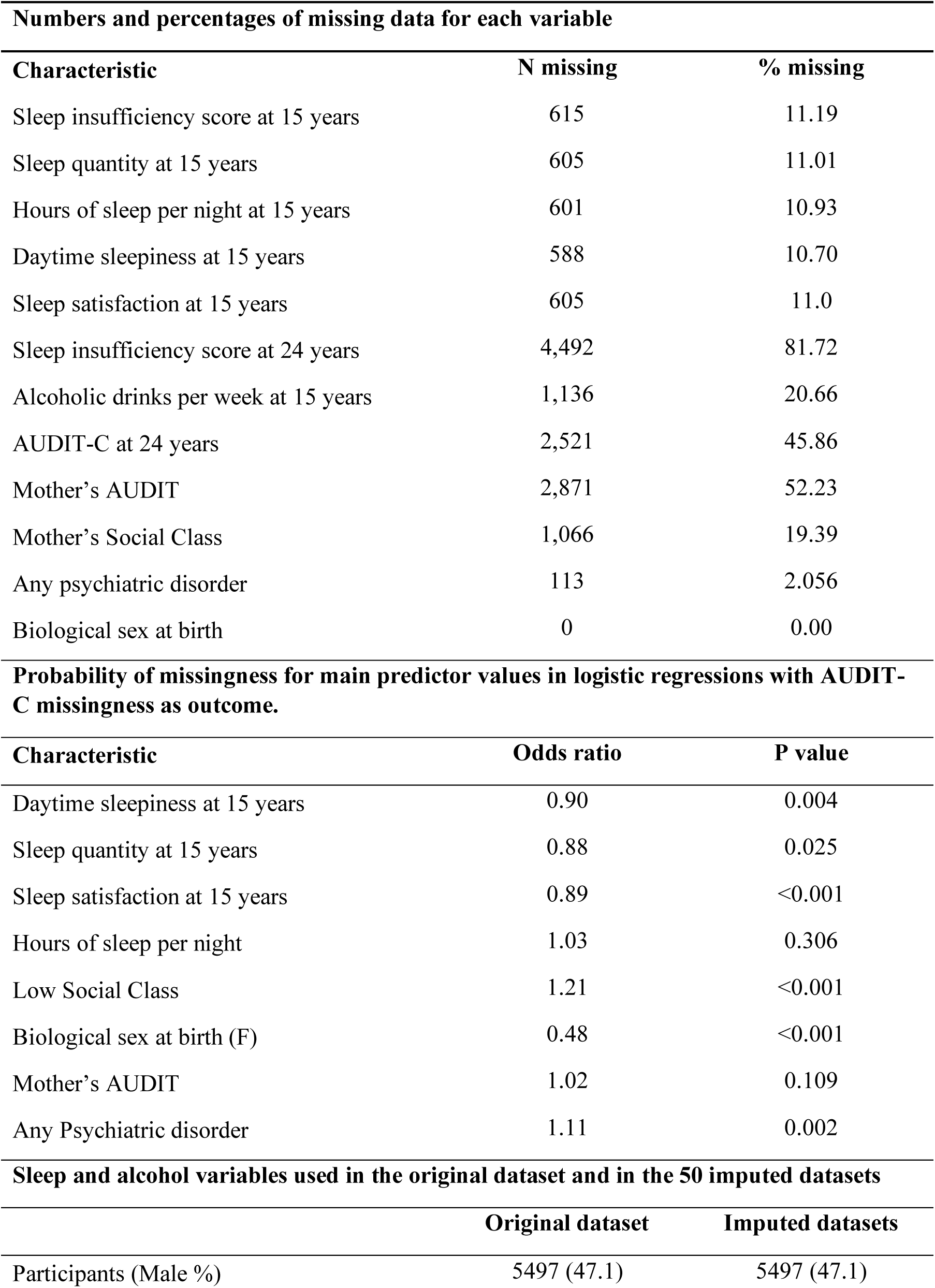
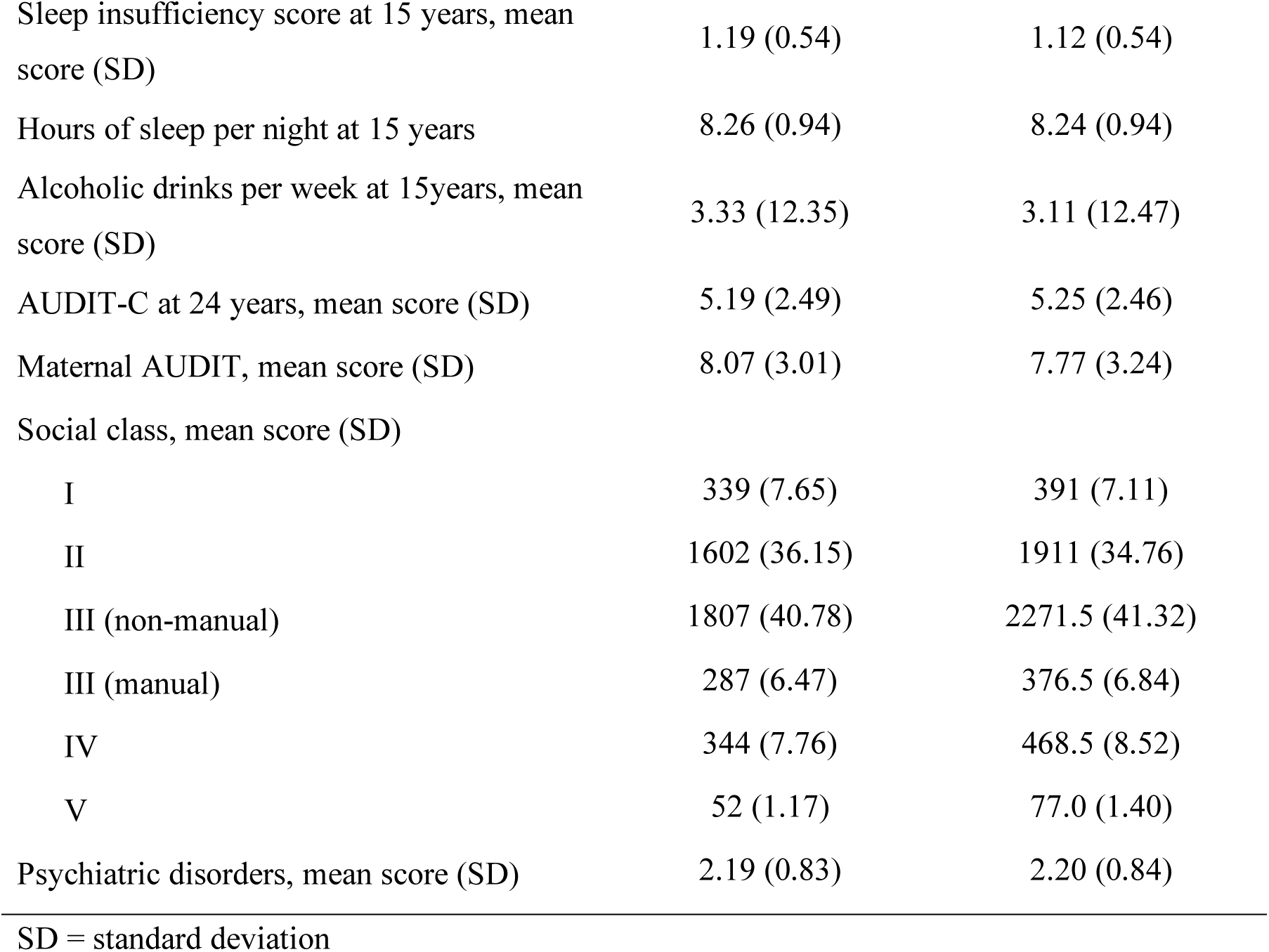
Missing values, population, sleep, and alcohol variables.

| **Numbers and percentages of missing data for each variable** |  |  |
| --- | --- | --- |
| **Characteristic** | **N missing** | **% missing** |
| Sleep insufficiency score at 15 years | 615 | 11.19 |
| Sleep quantity at 15 years | 605 | 11.01 |
| Hours of sleep per night at 15 years | 601 | 10.93 |
| Daytime sleepiness at 15 years | 588 | 10.70 |
| Sleep satisfaction at 15 years | 605 | 11.0 |
| Sleep insufficiency score at 24 years | 4,492 | 81.72 |
| Alcoholic drinks per week at 15 years | 1,136 | 20.66 |
| AUDIT-C at 24 years | 2,521 | 45.86 |
| Mother's AUDIT | 2,871 | 52.23 |
| Mother's Social Class | 1,066 | 19.39 |
| Any psychiatric disorder | 113 | 2.056 |
| Biological sex at birth | 0 | 0.00 |
| **Probability of missingness for main predictor values in logistic regressions with AUDIT-C missingness as outcome.** |  |  |
| **Characteristic** | **Odds ratio** | **P value** |
| Daytime sleepiness at 15 years | 0.90 | 0.004 |
| Sleep quantity at 15 years | 0.88 | 0.025 |
| Sleep satisfaction at 15 years | 0.89 | <0.001 |
| Hours of sleep per night | 1.03 | 0.306 |
| Low Social Class | 1.21 | <0.001 |
| Biological sex at birth (F) | 0.48 | <0.001 |
| Mother's AUDIT | 1.02 | 0.109 |
| Any Psychiatric disorder | 1.11 | 0.002 |
| **Sleep and alcohol variables used in the original dataset and in the 50 imputed datasets** |  |  |
|  | **Original dataset** | **Imputed datasets** |
| Participants (Male %) | 5497 (47.1) | 5497 (47.1) |
| Sleep insufficiency score at 15 years, mean score (SD) | 1.19 (0.54) | 1.12 (0.54) |
| Hours of sleep per night at 15 years | 8.26 (0.94) | 8.24 (0.94) |
| Alcoholic drinks per week at 15years, mean score (SD) | 3.33 (12.35) | 3.11 (12.47) |
| AUDIT-C at 24 years, mean score (SD) | 5.19 (2.49) | 5.25 (2.46) |
| Maternal AUDIT, mean score (SD) | 8.07 (3.01) | 7.77 (3.24) |
| Social class, mean score (SD) |  |  |
| I | 339 (7.65) | 391 (7.11) |
| II | 1602 (36.15) | 1911 (34.76) |
| III (non-manual) | 1807 (40.78) | 2271.5 (41.32) |
| III (manual) | 287 (6.47) | 376.5 (6.84) |
| IV | 344 (7.76) | 468.5 (8.52) |
| V | 52 (1.17) | 77.0 (1.40) |
| Psychiatric disorders, mean score (SD) | 2.19 (0.83) | 2.20 (0.84) |

SD = standard deviation

#### Statistical analysis

Continuous variables were described by mean and standard deviation (SD). Categorical data were expressed as percentages. To determine whether reduced sleep quantity and quality, daytime sleepiness, and hours of sleep were independent risk factors for alcohol use, we used multivariable linear regression with adjustment for the covariates. Separate models were used to evaluate the association between sleep variables and alcohol consumption at 15 years, the association between sleep variables and AUDIT-C at 24 years, and the correlation between the sleep insufficiency score at 15 years and AUDIT-C score at 24 years (longitudinal model). The longitudinal model was also adjusted for alcohol consumption at 15 years to evaluate the long-term effect of insufficient sleep accounting for long course alcohol consumption, thus including an auto-regressive component.

We examined model diagnostics using residual plots versus predicted plots and quantile–quantile plots. All model assumptions were met adequately.

We also conducted sensitivity analyses that were restricted to individuals who had data in both the exposure and outcome variables (complete cases analysis). Results of analyses based on each imputed dataset were combined using Rubin’s rules [49]. All analyses were conducted using R software, version 4.0.1 (R Core Team, 2020).

To further investigate the directionality of the associations we constructed a cross-lagged autoregressive path model [50] using the sleep insufficiency scores at 15 and 24 years, and alcohol measures at 15 and 24 years (the minmax scaled number of drinks per week and the minmax scaled AUDIT-C score, respectively). Measurement invariance of the alcohol measures and sleep insufficiency constructs at the two time points were verified prior to analyzing the cross-lagged panel model. To evaluate whether sleep insufficiency predicted alcohol consumption and AUDIT-C score, alcohol measures were used as dependent variables, with the sleep insufficiency score as independent variable and vice versa for the reverse lagged association. Model paths were estimated using the unweighted least squares method with confidence interval bootstrapping while controlling for the three covariates that resulted as significant predictors of alcohol use in the longitudinal regression analysis (i.e., sex, maternal AUDIT-C and social class). The following predetermined parameters were chosen to evaluate adequate model fit: root mean square error of approximation below 0.08 and a comparative fit index and Tucker-Lewis Index of 0.95 or higher [51]. Significant pathways were established based on α = .05. As for the other analyses we applied the CLPM to the imputed datasets and to the original dataset as sensitivity analysis. The CLPM were estimated using the lavaan R package version 0.6-12 (R Project for Statistical Computing).

### Preclinical study

#### Animals

Marchigian Sardinian alcohol preferring (msP) rats are characterized by high preference and high alcohol consumption levels, which make them a good model to study voluntary alcohol drinking and AUD since a young age [52–54]. Male rats were kept under a 12-hour dark/light cycle, constant temperature with water and food *ad libitum*. All animal care and experimental procedures were carried out in accordance with the guidelines of the European Community Council Directive for Care and Use of Laboratory Animals (2010/63/EU), Decreto Legislativo Italiano 26 (2014), and authorized by the local Animal Care Committee and by the Italian Ministry of Health.

#### Chronic sleep restriction and alcohol exposure

We attempted to model adolescent patterns of chronic sleep deprivation on weekdays and sleep compensation combined with alcohol exposure on weekends, adapting previous protocols of chronic sleep restriction and 2-bottle choice alcohol exposure [55, 56]. At post-natal day (P) 25, rats were divided into two groups: the chronic sleep restricted (CSR) group and the yoke controls (YC). The CSR group (n=22) was allowed to sleep for 4 hours every day during the first 4 hours of the light period, and sleep restricted for the remaining 20 hours, for four consecutive days. Rats were housed in pairs to avoid the confounding effect of social isolation, and chronic sleep restriction was achieved by using sleep deprivation cages (AM Microsystem, Urbisaglia, Italy) equipped with a programmable rotating bar swiping the cage floor at 2-3 rpm and changing direction at random intervals between 20 sec to 2 minutes. To control for the stress induced by the bar rotation, the YC group (n=18) was subjected to bar rotation for 6 hours during the dark period (1 hour after lights off) and left undisturbed for the remaining 18 hours. On the fifth day, both groups were left undisturbed. This cycle of 4 days of sleep restriction plus 1 day of recovery was repeated 6 times across the entire adolescence, which, in rats, spans from about P25 to P55 [57, 58].

Rats were exposed to a two-bottle choice (water and 10% alcohol) at P23 and P25 to determine baseline levels of alcohol preference and intake, and then on the recovery day throughout the 6 cycles of the CSR procedure (at P30, P35, P40, P45, P50, P55), and on alternating days between P90 and P96. Alcohol was available for 10 hours during the dark period, starting one hour after the light offset. The positions of the bottles were alternated. To measure alcohol and water intake, on the recovery day, a Plexiglas panel was inserted in the middle of the cage to temporarily separate the two rats. Measures of drinking and alcohol intake were recorded 30 minutes and 10 hours after bottle presentation.

#### EEG/EMG recordings and analysis

Under isoflurane anesthesia (1.5-2% volume) 9 rats (P38-50) were implanted bilaterally with epidural screw electrodes over the frontal (AP: +2 mm, ML: ±2 mm) and parietal (AP: - 2mm, ML: ±3mm) cortices, and the cerebellum as reference and ground for chronic electroencephalographic (EEG) recordings. Electrodes were fixed to the skull with dental cement. Two stainless steel wires were inserted into neck muscles to record electromyogram (EMG).

After 7 days of recovery, rats were transferred to the sleep restriction chambers, and after 24 hours of habituation, connected to the acquisition board (Open Ephys) by a recording head stage (Intan Technologies) and a commutator. Polysomnographic recordings were carried out continuously for 7 days and consisted of 24 hours of undisturbed baseline, 4 days of bar rotation (20 hours/day for the CSR group, 6 hours/day for the YC group), and 2 days of undisturbed recovery. EEG and EMG signals were filtered (EEG: high-pass filter at 0.1 Hz; low-pass filter at 100 Hz; EMG: high-pass filter at 10 Hz; low-pass filter at 70 Hz). All signals were sampled at 1000 Hz and down sampled at 512Hz for analysis. Waking, NREM sleep, and REM sleep were manually scored off-line (SleepSign, Kissei COMTEC, Matsumoto, Japan) in 4-s epochs according to standard criteria.

#### Behavioural tests

The first testing session started about 36 hours after the last ethanol exposure cycle (P57), whereas the second session started after 5 weeks of recovery (> P105). All tests were performed during the dark phase.

##### Open field test

Rats were habituated to the test room for 10 minutes and then placed at the edge of a white Plexiglas arena (43.4 x 43.4 x 30.3 cm) and allowed to move freely for 20 mins while being recorded (Med Associates, St Albans, VT, USA).

##### Light dark box

A Plexiglas box (40 x 40 x 40cm) was divided into two compartments connected by a sliding door (10 cm × 10 cm). The light chamber was brightly illuminated by a lamp. Animals were allowed to habituate to the test room for 5 mins and then placed in the centre of the light chamber facing the wall of the arena to explore the two chambers. The test lasted 5 mins.

##### Novelty suppressed feeding test

After 24hrs of food deprivation, each animal was placed in a novel Plexiglas arena (L 76.5 cm x 76.5 cm x h 40 cm) with a brightly illuminated center (∼1000Lux) containing a previously weighed food pellet. The animals were allowed free exploration and access to the food pellet for 5 minutes: the latency to their first bite and the amount of food consumed were measured. Rats that did not consume any food were automatically assigned a latency of 300 sec. After the 5 minutes, rats were returned to their home cage and allowed free access to another food pellet for 5 minutes to compare feeding behavior in a familiar environment.

##### Forced swim test

Rats were placed in a cylindrical container (50cm height and 20cm in diameter) filled with water (25±2°C) to a depth of 30cm for 6mins. Only activities in the final 4mins were considered for analysis. Behaviors were manually scored by two researchers blind to the animal’s condition. Immobility was considered as when the limbs and shrunk were immobile only to allow for floating.

#### Statistical analysis

Statistical tests used to analyze EEG, alcohol drinking, and behavioral data included ANOVA, Mann-Whitney, and Linear Mixed effect models, as detailed in the figure legends. Post-hoc tests included Dunnett’s multiple comparisons against baseline for EEG data and Šídák’s multiple comparisons test for alcohol drinking and behavioral measures. Bivariate correlations between behavioral tests data were evaluated using non-parametric Spearman’s test. The significance level alpha was set to 0.05. Analysis was performed with GraphPad Prism 9.5.1 and Python 3.9.

## Results

In the first part of this study, we carried out cross-sectional and longitudinal analyses of sleep insufficiency and alcohol use data collected over an interval of 9 years. Information on the population, sleep and alcohol variables in the imputed and complete dataset is summarized in Table 1. All models were adjusted for the presence of psychiatric disorders, maternal AUDIT-C and social class, and biological sex at birth.

### Cross-sectional effect of sleep deprivation on alcohol consumption at 15 and 24 years of age

Shorter and low-quality sleep, defined by both distinct sleep variables as well as the multidimensional score of sleep insufficiency, were associated with higher alcohol drinking at 15 years of age (Table 2). For sensitivity analyses, the same models were tested in the complete cases dataset in which we confirmed the association between the insufficient sleep score and alcohol consumption (Table 3). Conversely, we did not find any association between sleep and AUDIT-C score at 24 years.

**Table 2.** Cross-sectional and longitudinal associations between sleep parameters and alcohol consumption.

| **Cross-sectional association between sleep parameters and alcohol consumption at 15 years of age.** |  |  |  |
| --- | --- | --- | --- |
| **Parameter** | **Estimate** | **95% CI<sup>1</sup>** | **p-value** |
| Sleep insufficiency score | 1.7 | 0.84, 2.6 | <0.001 |
| Hours of sleep per night | -0.92 | -1.5, -0.38 | 0.001 |
| Daytime sleepiness | 1.6 | 0.71, 2.5 | <0.001 |
| Sleep quantity * | 0.99 | 0.16, 1.8 | 0.020 |
| Sleep satisfaction * | 0.65 | 0.09, 1.2 | 0.023 |
| **Longitudinal association between sleep parameters at 15 years and AUDIT-C at 24 years** |  |  |  |
| **Parameter** | **Estimate** | **95% CI<sup>1</sup>** | **p-value** |
| Hours of sleep per night | -0.11 | -0.20, -0.03 | 0.009 |
| Sleep insufficiency score | 0.35 | 0.21, 0.5 | <0.001 |
| Daytime sleepiness | 0.16 | 0.05, 0.26 | 0.003 |
| Sleep quantity * | 0.25 | 0.09, 0.40 | 0.002 |
| Sleep satisfaction * | 0.24 | 0.15, 0.34 | <0.001 |
| **Evaluation of autoregressive, concurrent, and cross-lagged correlations between insufficient sleep score and alcohol use.** |  |  |  |
| **Autoregressive correlation** | **Estimate** | **95% CI<sup>1</sup>** | **p-value** |
| Alcohol T1 → Alcohol T2 | 0.01 | -0.095 - 0.11 | 0.88 |
| **Cross lagged correlation** | **Estimate** | **95% CI<sup>1</sup>** | **p-value** |
| Sleep T1 → Alcohol T2 | **0.021** | **0.012 - 0.03** | **<0.001** |
| **Concurrent correlation** | **Estimate** | **95% CI<sup>1</sup>** | **p-value** |
| Sleep T1 → Alcohol T1 | **0.0037** | **0.001 - 0.006** | **0.039** |

\* Responses are on an inverse scale, with higher scores indicating less sleep or low satisfaction. <sup>1</sup> CI = Confidence Interval. Sleep T1=insufficient sleep score at 15 years of age. Alcohol T1=alcohol consumption at 15 years of age. Alcohol T2=AUDIT-C score at 24 years of age.

**Table 3.** Evaluation of the cross-sectional and longitudinal associations in the complete case analysis.

| **Cross-sectional association between sleep insufficiency score and alcohol consumption at 15 years of age in the complete case analysis** |  |  |  |
| --- | --- | --- | --- |
| **Characteristic** | **Estimate** | **95% CI<sup>1</sup>** | **p-value** |
| **Sleep insufficiency score** | **0.89** | **0.30, 1.5** | **0.003** |
| Biological sex at birth (F) | -0.53 | -1.1, 0.08 | 0.088 |
| Any psychiatric disorders | 1.1 | 0.71, 1.5 | <0.001 |
| Mother's AUDIT | 0.10 | -0.01, 0.20 | 0.066 |
| Mother's social class | 0.27 | -2.0, 2.6 | 0.8 |
| **Longitudinal association between sleep insufficiency score at 15 years of age and AUDIT-C score at 24 years of age in the complete cases analysis** |  |  |  |
| **Characteristic** | **Beta** | **95% CI<sup>1</sup>** | **p-value** |
| **Sleep insufficiency score** | **0.26** | **0.00, 0.52** | **0.048** |
| Alcohol consumption at 15 years | 0.05 | 0.02, 0.07 | 0.001 |
| Biological sex at birth (F) | -0.83 | -1.1, -0.56 | <0.001 |
| Any psychiatric disorders | 0.01 | -0.17, 0.19 | >0.9 |
| Mother's AUDIT | 0.10 | 0.05, 0.14 | <0.001 |
| Mother's social class | -0.80 | -2.0, 0.43 | 0.2 |

<sup>1</sup>CI = Confidence Interval

### Longitudinal effect of sleep deprivation on AUDIT-C score

We then tested whether insufficient sleep at 15 years predicted AUDIT-C score at 24 years using a linear model that included an autoregressive term for alcohol consumption at 15 years. We found that higher alcohol consumption at 24 years was associated with fewer hours of sleep during weekdays, lower sleep quantity and satisfaction, a higher daytime sleepiness, and a higher insufficient sleep score at 15 years (Tables 2 and 3).

To assess the directionality of the longitudinal association, we used an autoregressive cross-lagged path model and found that sleep insufficiency at 15 years had a significant effect on AUDIT-C at 24 (Sleep T1 → Alcohol T2). Conversely, the estimated number of alcoholic drinks per week at 15 years of age did not predict sleep insufficiency at 24 years (Alcohol T1 → Sleep T2) (Figure 1).

**Figure 1.**
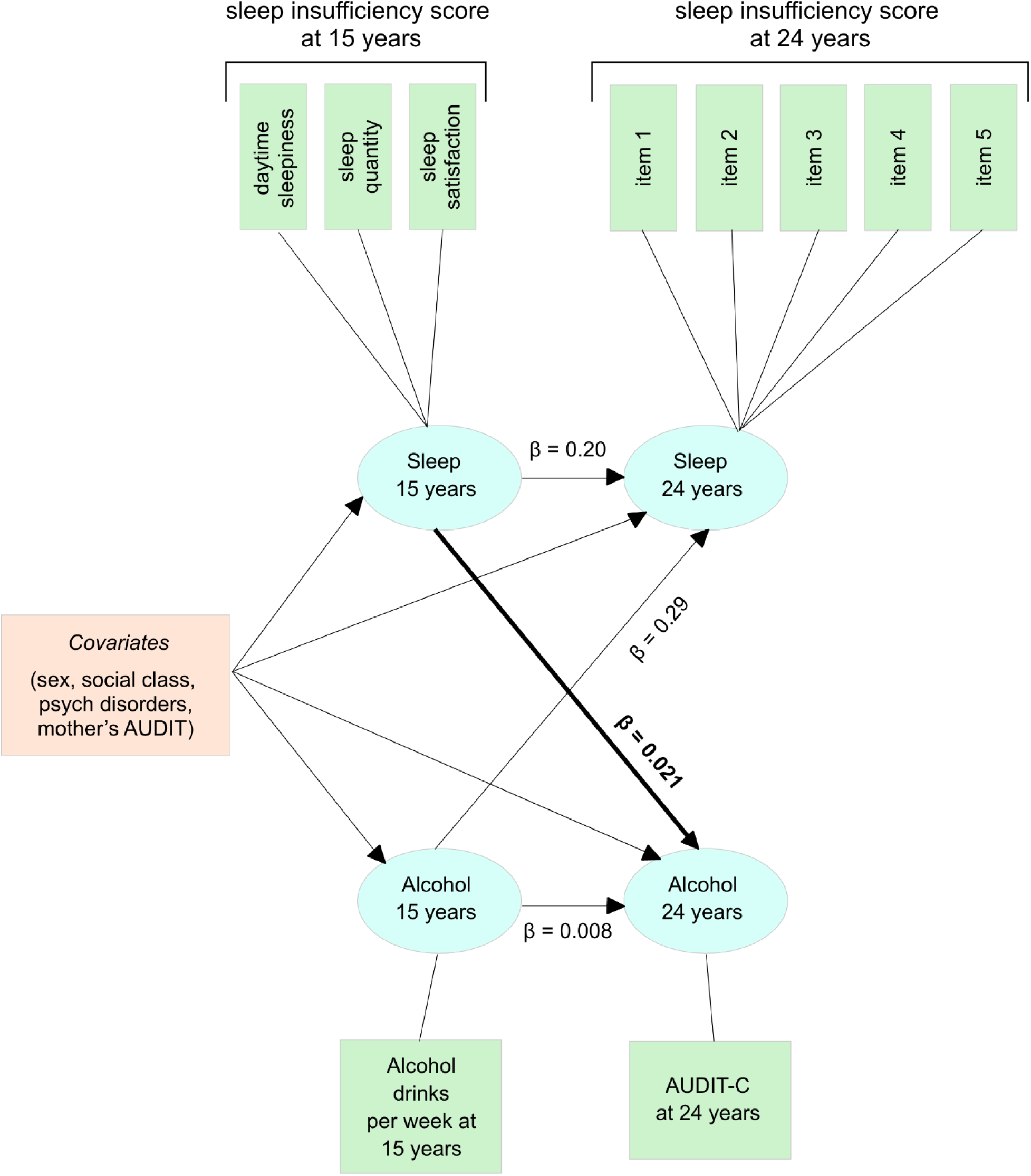
Cross-lagged panel model evaluating the cross-sectional and longitudinal association between sleep and alcohol. Significant paths and relative estimates are in bold.

In summary, these results support the hypothesis of a unidirectional causal relation between sleep insufficiency in mid-adolescence and alcohol drinking in emerging adulthood.

### Chronic sleep restriction escalates alcohol drinking during adolescence and adulthood in msP rats

To model and determine the impact of adolescent patterns of sleep deprivation in a controlled preclinical experiment, we subjected adolescent msP rats to six cycles of 4-day sleep restriction followed by 1 day of recovery sleep, throughout their adolescence (Figure 2A). The automated sleep restriction protocol was validated in a cohort of msP rats using polysomnographic recordings and showed a reduction in total time spent in NREM and REM sleep on average of 53% and 47%, respectively, and an increase of 43% in total wake duration relative to 24-hour of undisturbed baseline (Figure 2B). In controls, there was no significant effect on the time spent in NREM and REM sleep relative to the 24-hour baseline (Figure 2C).

**Figure 2.**
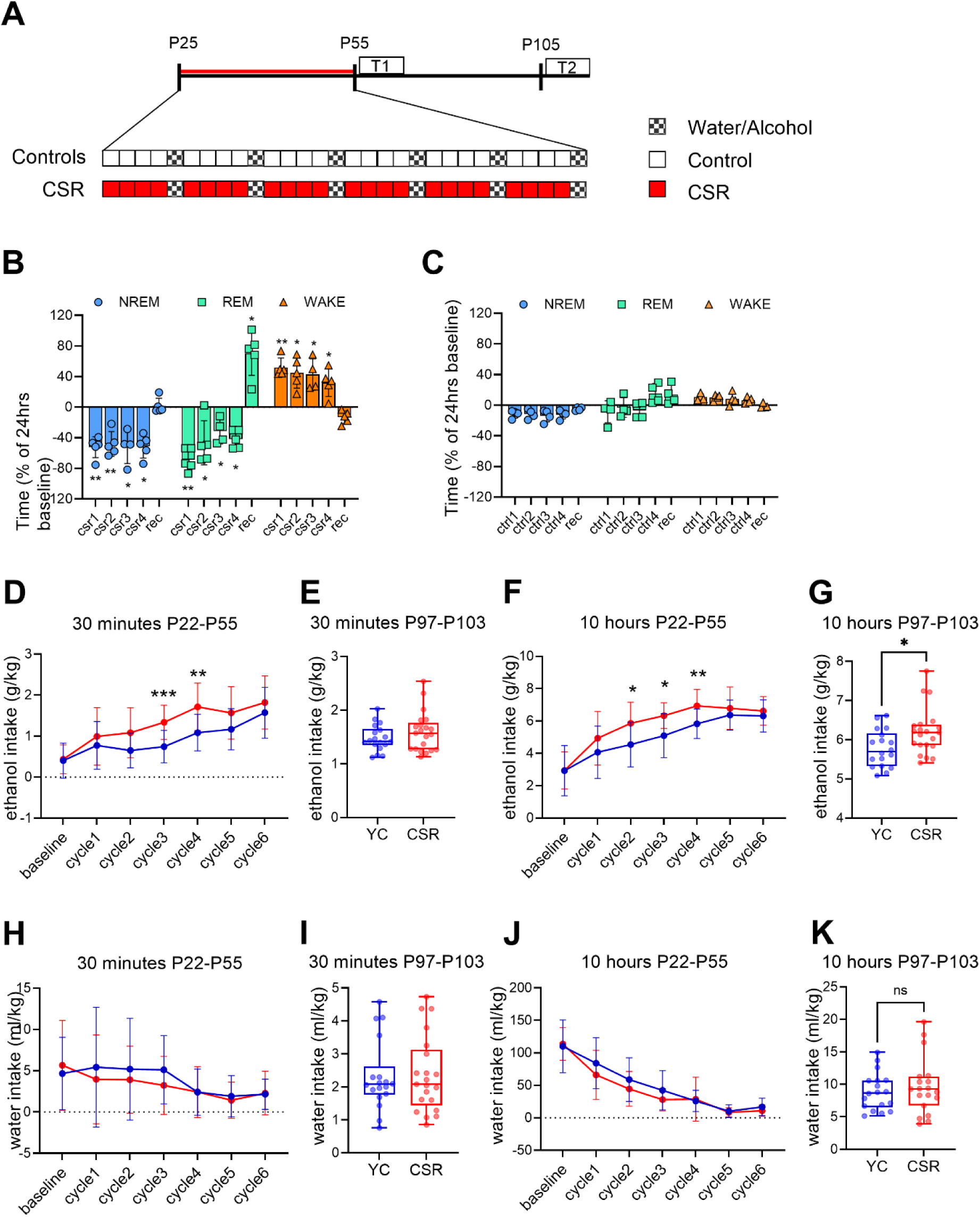
Experimental design, polysomnography, and alcohol and water consumption. **A)** Experimental design for adolescent chronic sleep restriction (CSR) and yoked control (YC) groups. Each white rectangle represents 1 day of 6-hour bar rotation (Ctrl); each red rectangle represents 1 day of 20-hour bar rotation (CSR); yellow-gray rectangles represent recovery days during which rats could drink water and ethanol. CSR = chronic sleep restriction; YC = yoked controls; Ctrl = control; T1 and T2 indicate batteries of behavioral tests. **B)** Time spent in each physiological state, expressed as % of a 24-hour baseline, for the CSR group. Repeated measure ANOVA was performed on raw data (minutes), for NREM sleep F_day_(1.789, 7.155)=13.11 p=0.0045; for REM sleep, F_day_(2.23,10.25)=39.35 p<0.0001; for wake F_day_(1.57, 7.21)=18.29 p=0.002. Dunnett’s multiple comparisons post hoc tests against baseline. **C)** Time spent in each physiological state, expressed as % of a 24-hour baseline, for the YC group. CSR = chronic sleep restriction; YC = yoked controls; Ctrl = control; rec = recovery. * p<0.05, ** p<0.01, *** p<0.001, **** p<0.0001. **D)** alcohol intake across the 6 cycles of sleep restriction, measured 30 minutes after presenting the 2 bottles (water and 10% ethanol). LME model F_group_(1,38)=16.24 p=0.0003. Post hoc Šídák’s multiple comparisons test, cycle 3 adjusted p=0.0003, cycle 4 p=0.0027. **E**) average ethanol consumption, measured 30 minutes after bottle presentation, over 7 days of intermittent (1 day on, 1 day off) 2-bottle choice in adulthood, Mann Whitney test p=0.46. **F)** alcohol intake across the 6 cycles of sleep restriction, measured 10 hours after presenting the 2 bottles (water and 10% ethanol). LME model F_group_(1,36)=15.59 p=0.0004. Post hoc Šídák’s multiple comparisons test, adjusted p value for cycle 2 p=0.0326, cycle 3 p=0.0155, and cycle 4 p=0.0078. **G)** average ethanol consumption, measured 10 hours after bottle presentation, over 7 days of intermittent (1 day on, 1 day off) 2-bottle choice in adulthood, Mann Whitney test p=0.03. **H)** water intake across the 6 cycles of sleep restriction, measured 30 minutes after presenting the bottles. LME model F_group_(1,38)=0.55, p=0.46. **I)** water intake across the 6 cycles of sleep restriction, measured 10 hours after presenting the bottles. LME model F_group_(1,38)=1.95, p=0.17. **J)** and **K)** average water consumption, measured 30 minutes (J) and 10 hours (K) after bottle presentation, over 7 days of intermittent (1 day on, 1 day off) 2-bottle choice in adulthood, Mann Whitney test p=0.91 and p=0.77. CSR group in red (N=22), YC group in blue (N=18). ns= not significant, * p<0.05, ** p<0.01, *** p<0.001.

Both groups showed a gradual increase in alcohol consumption across adolescence when exposed to an intermittent 2-bottle choice paradigm during the recovery day. However, rats that underwent CSR showed a faster escalation in alcohol drinking relative to controls, reaching higher levels of alcohol intake at earlier times (Figure 2D, F). When rats were tested again for alcohol intake after 5 weeks of rest, increased alcohol consumption persisted in the CSR group relative to controls (Figure 2 E, G). Water consumption remained similar between groups across the entire experiment (Figure 2H-K).

### Sleep restriction and alcohol drinking promote short- and long-lasting risky behavior in anxiogenic environment

To identify whether CSR and alcohol drinking were associated with short and/or long-lasting behavioral changes, we performed two batteries of tests (Figure 2A).

In the open field (OF) test, while the controls showed a significant decline in time spent moving in the arena and in the distance traveled from late adolescence to young adulthood, rats previously exposed to CSR maintained higher, adolescent-like values of travelling when tested later in life (Figure 3A, left). Conversely, the two groups spent a similar amount of time in the center of the arena relative to the edges (Figure 3A, right). A subgroup of rats underwent the light/dark box test, where CSR rats showed a tendency to spend more time in the lit box and to enter the lit chamber relative more rapidly than controls (Figure 3B).

**Figure 3.**
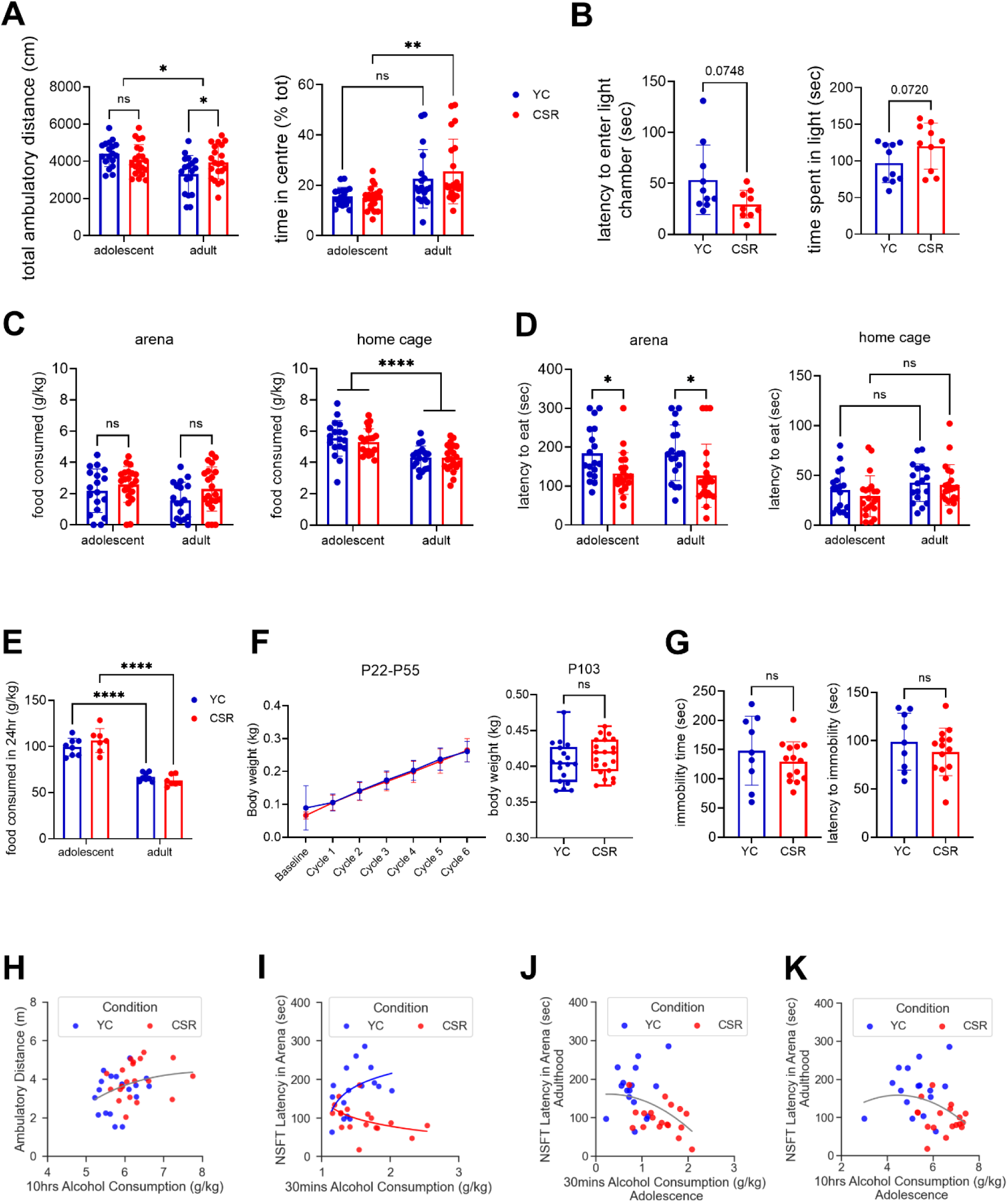
Behavioral tests. **A)** Open field test. *Left*, open field test shows a decline in total distance traveled from adolescence to adulthood in the YC group (N=18) but not in the CSR group (N=22). Two-way ANOVA found a main effect of age F_age_(1,37)=6.976 p=0.012, and an interaction between age and group F_group_ x _age_(1,37)=4.35 p=0.043, which were confirmed by Šídák’s multiple comparisons post hoc tests, adolescent p=0.43, adult p=0.049, YC adolescent vs adult p=0.0053, CSR adolescent vs adult p=0.9. *Right,* time spent in the center of the open field. Two-way ANOVA found only an effect of age F_age_(1,37)=17.30 p=0.0002, confirmed by Šídák’s multiple comparisons post hoc tests: YC p=0.057, CSR p=0.0015. **B)** Light-dark box. We found a trend towards shorter latency to enter the light box and longer time spent in the light box in CSR (N=9) relative to YC (N=10). p value for Mann-Whitney test displayed. **C)** Novelty Suppressed Feeding test. *Left,* food consumption in the arena, two-way repeated measure (rm) ANOVA F_group_(1,37)=3.16 p=0.083, F_age_(1,37)=3.04 p=0.089, F_age x group_(1,37)=0.36 p=0.54. *Right,* food consumption in home-cage, two-way rmANOVA F_age_(1,37)=56.37 p<0.0001, F_group_(1,37)=0.2 p=0.6, F_age x group_(1,37)=0.55 p=0.46, Šídák’s post hoc test for YC and CSR p<0.0001. **D)** Novelty Suppressed Feeding test. *Left,* latency to eat in the arena, two-way rmANOVA F_group_(1,37)=13.35 p=0.0008, F_age_(1,37)=0.01 p=0.9, F_age x group_(1,37)=0.05 p=0.8. Šídák’s multiple comparisons post hoc tests, adolescent p=0.047, adult p=0.018. Right, latency to eat in the home cage, two-way rmANOVA F_group_(1,37)=0.66 p=0.42, F_age_(1,37)=5.74 p=0.022, F_age x group_(1,37)=0.22 p=0.64. **E)** food consumption over a period of 24 hour, two-way rmANOVA F_group_(1,13)=0.18 p=0.67, F_age_(1,13)=165,9 p<0.0001, F_age x group_(1,13)=3.5 p=0.083, Šídák’s post hoc test for YC (N=8) and CSR (N=7) p<0.0001. **F)** Measures of body weight during adolescence (left) and young adulthood (right), CSR N=22, YC N=18, Mann-Whitney p=0.22. **G)** Forced swim test. Right, time spent immobile, Mann-Whitney p=0.43. Left, latency to immobility Mann-Whitney p=0.48. YC= yoke controls; CSR= chronic sleep restriction; NSFT= novelty suppressed feeding test; ns= not significant, * p<0.05, ** p<0.01, ****p<0.0001. **H** Positive correlation between alcohol consumption and ambulatory distance in adult rats (N total=39; ρ=0.37 p=0.022). **I)** Correlation between the amount of alcohol consumed at 30 minutes and the latency to approach food in the arena in CSR adult rats (N=18; ρ=-0.59, p=0.01) and YC (N=16; ρ=0.49 p=0.054). **J)** and **K)** Longitudinal correlation between adolescent alcohol consumption and adult performance in the novelty suppressed feeding test (NSFT) (N= 34; ρ=-0.45 p=0.008 at 30 minutes *left*, ρ=-0.44 p=0.01 at 10 hours *right*). ρ= Spearman’s coefficient. For correlations with the latencies at the NSFT, rats that did not engage in food consumption during the test were removed from the analysis. YC=yoke controls; CSR = chronic sleep restriction.

In the novelty suppressed feeding (NSF) test, we found that the amount of food consumed was unchanged between controls and CSR, both in the arena and in the home cage (Figure 3C). However, CSR rats showed reduced latency to approach and eat food in the unfamiliar arena, whereas in the home cage no significant difference was detected (Figure 3D). The shorter latency was maintained when animals were tested again after 5 weeks of recovery. To rule out the possibility that the decreased latency was due to an increased appetite in response to the CSR procedure, we measured food consumption during the recovery period and found no difference between the groups (Figure 3E). Moreover, 6 cycles of CSR affected neither growth rate nor body weight throughout the experiment (Figure 3F). No differences were found between groups in the forced swim test (FST, Figure 3G).

In summary, adult rats exposed to adolescent CSR showed increased locomotion and decreased signs of anxiety relative to controls. No alterations in despair levels were highlighted.

We then explored possible correlations between behavioral parameters and alcohol consumption and found that, in adult rats, alcohol consumption positively correlated with OF ambulatory time and distance (Figure 3H), and with time spent in the center of the OF arena. In adolescence, CSR rats with the highest levels of alcohol consumption in the first 30 minutes were also the fastest to approach the food in the NSF test arena (Figure 3I). By contrast, controls showed an opposite trend. Finally, we found that the average amount of alcohol consumed during adolescence negatively correlated with the latency of approaching food in the NSF test arena later in adulthood (Figure 3J, K), highlighting a longitudinal link between adolescent alcohol consumption levels and increased risk-taking behavior later in life.

## Discussion

The study tested the hypothesis that insufficient sleep during adolescence promotes alcohol drinking in young adulthood by using both epidemiological and preclinical data. The epidemiological data involved a well-characterized cohort with self-reported sleep patterns and alcohol consumption data obtained 9 years apart. We found a cross sectional association between sleep and alcohol consumption at 15 years of age, with a 1.7 increase in weekly drinks for each unitary increase in the sleep insufficiency score. The autoregressive cross-lagged path model showed that insufficient sleep at 15 years predicted AUDIT-C scores at 24, emphasizing a prospective developmental effect over 9 years. These findings extend the results of other cross-sectional [18, 19, 59, 20] and longitudinal reports [21, 23, 24, 22, 26, 60–63] affirming the reproducibility of such associations across diverse population studies and extended timeframes.

However, no concurrent association was observed between sleep and AUDIT-C score at 24 years, and alcohol drinking at 15 years did not predict sleep problems at 24. These findings contrast with studies reporting bidirectional associations between insomnia and alcohol use in adolescents and adults [21, 64–67] and are unlikely explained by the high percentage of missing data for the sleep insufficiency score at 24 years (Table1) [68].

The enduring impact of sleep insufficiency on alcohol related behavior raises questions about the underlying reasons. Significant shortcomings intrinsic to clinical studies make it difficult to answer this question. Pre-existing altered developmental trajectories, peer influence and engagement in social activity might affect sleep and promote alcohol drinking without a causal relationship between the two [69]. Moreover, in our ALSPAC population, the sleep and alcohol measurements were self-reported and might have been affected by recall bias and social desirability bias.

To address this, we developed a preclinical rat model of adolescent sleep restriction and alcohol drinking. We found that if exposed to CSR across adolescence, genetically selected alcohol preferring msP rats escalated alcohol consumption and led to higher alcohol drinking in adulthood. Our experimental paradigm tried to control for the stress induced by the sleep restriction and assessed behavior over a long temporal interval. We validated the sleep restriction procedure by performing polysomnography in adolescent rats. Moreover, our sleep restriction protocol preserved a sleep opportunity window during the first hours of the light period, when rats are naturally more propense to be asleep and sleep pressure is higher [70]. This helped exclude the effect of a circadian shift in sleep, which can have additional effects on alcohol drinking behavior [71].

Notably, msP rats [52] are believed to drink alcohol partially as an anxiety relieving mechanism [54, 72, 73]. Indeed, control rats with the highest levels of alcohol consumption tended to be the most anxious when tested in the NSF arena. However, in CSR rats the opposite trend was observed and other typical markers of anxiety, such as the time spent in the center of the OF arena or the time spent in the light box, did not differ from controls. If anything, CSR rats showed less anxious-like behavior in all tests performed. At first glance, these results could appear in contrast with existent reports showing that repeated adolescent alcohol binge drinking [74] and sleep loss [75] can increase anxiety and depressive symptoms in early and mid-adulthood. On the other hand, it is also known that disrupted sleep is associated with global risky decision making in adolescents, including illicit substance use, sexual risk-taking, and road/transport-related safety [10]. Therefore, the elevated levels of alcohol consumption observed as a result of CSR can be attributed to an increased inclination toward risky behaviors, stemming from reduced anxiety levels.

In addition to heightened risk-taking, changes in emotional responses, altered regulation of reward processes [76, 77] and changes in preference for caloric beverages [78–81] may be also linked to sleep loss and increase alcohol consumption. While here we did not measure overall caloric intake, we attempted to control for this confounding factor by giving the rats *ab libitum* access to food throughout the experiment. 24-hour measurements of food consumption during the recovery period did not reveal a significant difference in overall food intake (g) between CSR and controls. Moreover, in our experimental paradigm, alcohol exposure occurred during the CSR recovery day, when theoretically, caloric intake should go down and be similar to control conditions [80]. Finally, increased alcohol intake persisted for many weeks of recovery after the sleep restriction procedure, when potential differences in energy expenditure should have ceased.

In conclusion, our findings favor the hypothesis that insufficient sleep during adolescence leads to higher alcohol consumption in adulthood and represents a contributing factor for future AUD associated with increased propensity to risk-taking behavior. Future experiments should explore the biological mechanisms linking sleep loss to the development of this complex neuropsychopathology and psychiatric comorbidity.

## Acknowledgments

We are extremely grateful to all the families who took part in this study, the midwives for their help in recruiting them, and the whole ALSPAC team, which includes interviewers, computer and laboratory technicians, clerical workers, research scientists, volunteers, managers, receptionists, and nurses. This publication is the work of the authors and LdV will serve as guarantor for the contents of this paper.

